# Viral gene drive in herpesviruses

**DOI:** 10.1101/717017

**Authors:** Marius Walter, Eric Verdin

## Abstract

Herpesviruses are ubiquitous pathogens in need of novel therapeutic solutions. Current engineered gene drive strategies rely on sexual reproduction, and are thought to be restricted to sexual organisms. Here, we report on the design of a novel gene drive system that allows the spread of an engineered trait in populations of DNA viruses and, in particular, herpesviruses. We describe the successful transmission of a gene drive sequence between distinct strains of human cytomegalovirus (human herpesvirus 5) and show that gene drive viruses can efficiently target and replace wildtype populations in cell culture experiments. Our results indicate that viral gene drives can be used to suppress a viral infection and may represent a novel therapeutic strategy against herpesviruses.

## Main text

Herpesviruses are universal pathogens that are implicated directly or indirectly in numerous human diseases. In particular, human cytomegalovirus (hCMV) is an important threat to immunocompromised patients, such as HIV-infected individuals, receivers of organ transplants, newborns and the elderly. While treatment options exist, drug resistance and adverse secondary effects render the development of new therapeutic solutions necessary.

Gene drive refers to the transmission of specific genetic sequences from one generation to the next with a high probability and can propagate a trait to an entire population (*1–7*). Most recent engineered gene drive strategies rely on CRISPR-Cas9 editing, where a Cas9 transgene is inserted in place of a natural sequence, alongside a guide RNA (gRNA) targeting the very same location. During sexual reproduction, repair of an unmodified allele by homologous recombination after cleavage by Cas9 leads to duplication of the synthetic sequence. This strategy relies on the simultaneous presence of a wildtype and a gene drive allele in the same cell nucleus. For these reasons, it has generally been assumed that an engineered gene drive could only be designed in sexually reproducing organisms, excluding bacteria and viruses (*5, 6*).

Herpesviruses are nuclear-replicating DNA viruses with large dsDNA genomes (100–200 kb) that encode 100 to 200 genes (*8*). They frequently undergo homologous recombination during their replication cycle and can be efficiently edited by CRISPR-Cas9 (*9–12*). These properties enabled the design of a new gene drive strategy that doesn’t involve sexual reproduction, but relies on coinfection of a given cell by a wildtype and an engineered virus (**Fig. 1A**). Upon coinfection, the wildtype genome is cut and repaired by homologous recombination, producing new gene drive viruses that progressively replace the wildtype population. Here we present a proof of concept for such a phenomenon, using hCMV as a model.

**Fig. 1.**
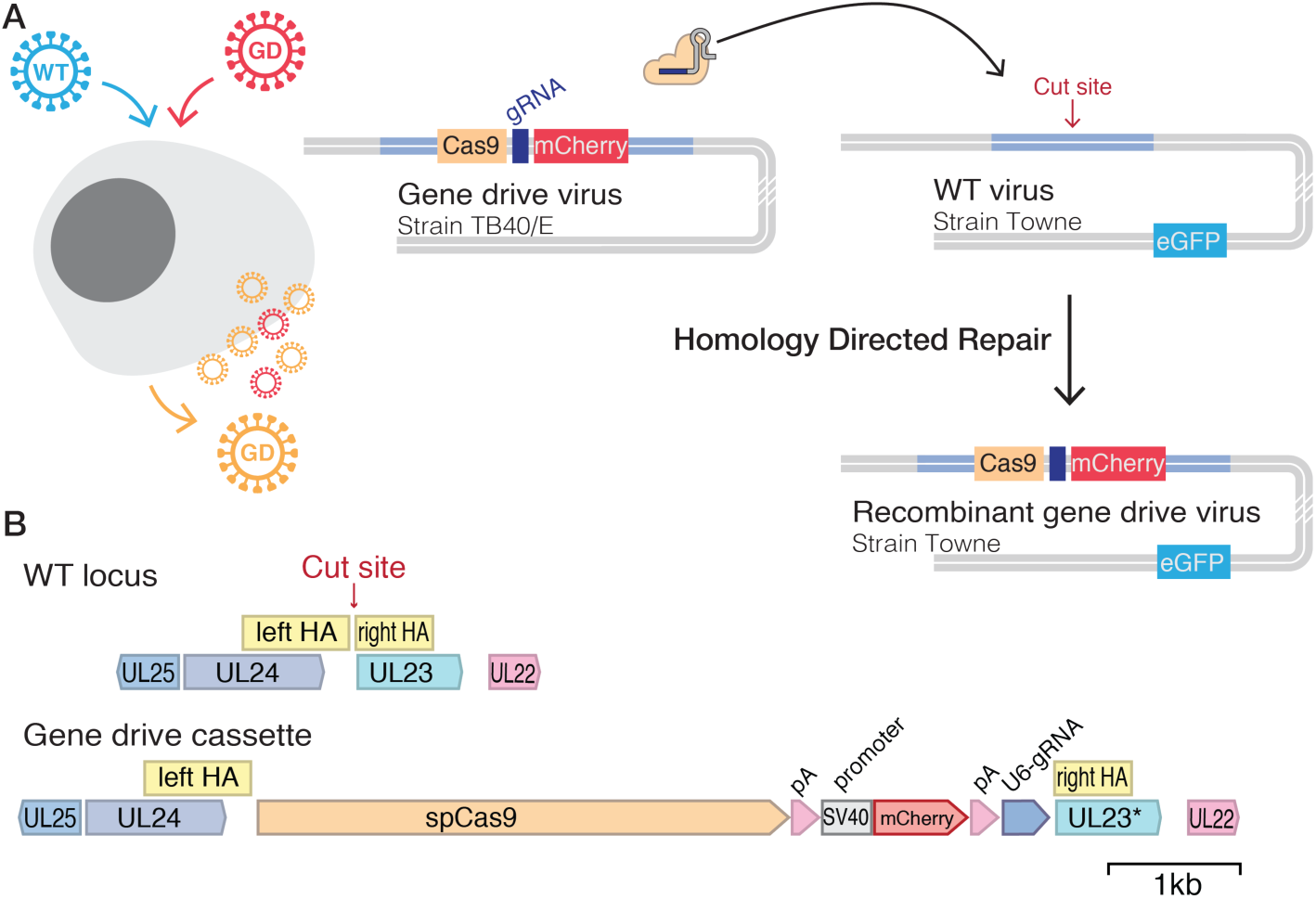
Design of a gene drive system in herpesviruses. **(A)** CRISPR-based gene drive sequences are, at a minimum, composed of Cas9 and a gRNA targeting the complementary wildtype (WT) locus and can harbor an additional ‘cargo’ that will be carried over with the rest of the sequence. When present in the same cell nucleus. Cas9 targets and cleaves the WT sequence. Homology-directed repair of the damaged WT locus using the gene drive sequence as a repair template ensures the conversion of the wildtype locus into a new gene drive sequence. In herpesviruses, gene drive involves the coinfection of a given cell by a WT and a modified virus. Cleavage and repair of the WT genome convert the virus into a new gene drive virus, spreading the modification into the viral population. In this example. hCMV WT virus (Towne strain) expresses eGFP florescent protein, and the gene drive virus (TB40/E strain) carries mCherry. Recombinant viruses then express both eGFP and mCherry. (**B**) Modified and unmodified *UL23* locus. The target site is located 13 bp upstream of the *UL23* coding sequence. Here, the gene drive cassette is composed of *Cas9*. the SV40 polyA signal. the SV40 promoter, an *mCherry* reporter. the beta globin polyA signa and a U6 driven gRNA.

We first aimed to build a gene drive system that would not affect viral infectivity, so that it could potentially spread easily into a wildtype population. We chose to insert a gene drive cassette into *UL23*, a viral gene that is dispensable for hCMV replication in human fibroblasts (*13*). We cloned a donor plasmid containing homology arms, *Cas9* (from *Streptococcus pyogenes)*, an *mCherry* fluorescent reporter and a gRNA targeting *UL23* 5′UTR (**Fig. 1B**). In this system, *Cas9* transcription is driven by the *UL23* endogenous viral promoter (**Fig. S1A**). Human foreskin fibroblasts were transfected with the gene drive plasmid and infected with TB40/E-bac4, a wildtype hCMV strain (*14*). mCherry-expressing viruses created by homologous recombination were isolated and purified until a pure population of gene drive virus (GD-mCherry) could be obtained (**Fig. S1B-C**). We similarly incorporated an *mCherry* reporter into a neutral region of TB40/E-bac4 (referred hereafter as simply TB40/E, **Fig. S1D**). GD-mCherry replicated with a slightly slower dynamic than TB40/E but ultimately reached similar titers (**Fig. S1E**).

To determine if the gene drive virus could recombine with unmodified virus, fibroblasts were co-infected with equal quantities of GD-mCherry and an eGFP-expressing wildtype virus from a different viral strain (Towne-eGFP) (*15*). The supernatant of co-infected cells was then used to infect fresh fibroblasts at a very low multiplicity of infection (MOI) (**Fig. S2A-B**). In this second generation, we detected cells and viral plaques expressing eGFP alone, mCherry alone, or mCherry and eGFP together (**Fig. 2A, Fig. S2C**). The presence of mCherry-eGFP-expressing viral plaques suggested that both mCherry and eGFP were inserted in the same viral genome.

**Fig. 2.**
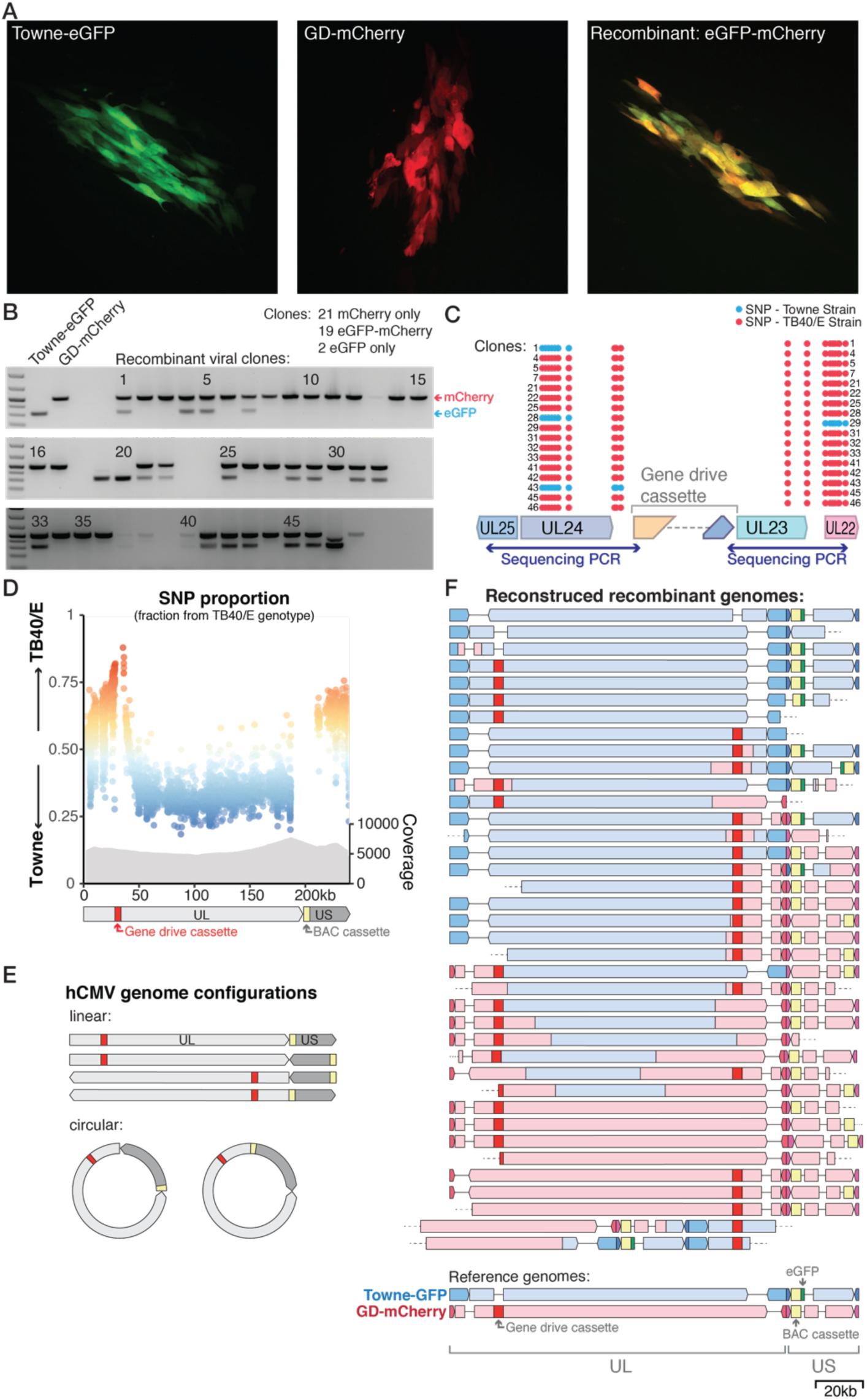
Recombination of the gene drive cassette into the wildtype genome. Fibroblasts were coinfected with Towne-eGFP (WT) and GD-mCherry, and supernatant was used to infect f resh cells. (**A**) Representative examples of fluorescent viral plaques spreading on fibroblasts, expressing either eGFP only (left), mCherry only (middle), or both (right). (**B**) PCR for mCherry (upper band) and eGFP (lower band) on 48 recombinant viral genomes. (**C**) PCR of homology arms and Sanger sequencing of 17 eGFP-mCherry expressing viral clones. Blue dots: SNPs from Towne strain; Red dots: TB40/E strain.(**D**) Fraction of SNPs of Towne or TB40/E origin alongside the hCMV genome. Each dot represents an individual SNP. Data combine two biological replicates after Oxford Nanopore sequenc ing. Coverage gives the number of reads, allowing multiple mapping. (**E**) The hCMV genome can be found in four linear or two circular genome configurations, depending on the respective orientation of long (UL) and short (US) domains. (**F**) Reconstruction of recombination history of individual hCMV genomes from long (>200 kb) Nanopore reads. One genome corresponds to one sequenc ing read and colors indicate the strain of origin. Gaps represent regions deleted compared to the reference genome, dashed lines indicate uncovered region.

To confirm this hypothesis, we first recovered episomal viral DNA from infected cells using a modified HIRT DNA extraction method (*16*). Both GD-mCherry and Towne-eGFP viral genomes carry BAC vector sequences that can be used to transform circular viral DNA into competent *E. coli* (*14, 15*). After transformation of episomal DNA from two independent coinfection experiments, 48 clones were isolated, each representing a distinct viral genome. Importantly, PCR revealed that 19 clones (40%) carried both *eGFP* and *mCherry*, whereas 21 clones (44%) were positive for *mCherry* only and only two clones (4%) for *eGFP* only (**Fig. 2B**). The presence of multiple *eGFP-mCherry* positive clones and the relative absence of *eGFP-*only suggested that most of Towne-eGFP viral genomes had incorporated the gene drive cassette containing *mCherry*. Towne and TB40/E viral strains are differentiated by numerous SNPs, allowing us to pinpoint homologous recombination breakpoints. Sanger sequencing and analysis of SNPs around the gene drive cassette showed that in 4 clones out of 17, homologous recombination had occurred immediately next to the CRISPR cut site (**Fig. 2C, Fig. S3**). This also highlighted that in most cases, homologous recombination breakpoint was located more that 1–2 kb away from the insertion site.

We therefore sought to use long-read sequencing to further investigate recombination between gene drive viruses (TB40/E strain) and unmodified viruses (Towne strain). Fibroblasts were co-infected (MOI=0.1 for both viruses) for 2 weeks, and virions collected from the culture supernatant. Linear viral DNA was extracted from purified virions and subjected to long-read sequencing using Oxford Nanopore sequencing. In two biological replicates, 41,779 and 9,926 reads with a length greater than 10 kb were recovered, of which 98.9% and 96.6%, respectively, could be mapped onto the hCMV genome (sequencing summary in **Fig. S4**). For each position along the hCMV genome, we plotted the proportion of SNPs originating from one strain or the other (**Fig. 2D, Fig. S5**). As expected, about 80% of SNPs immediately around the target site originated from the TB40/E donor strain, and this proportion decreased sharply further away from the cutting site. This finding indicated that the gene drive cassette was selectively integrated into the Towne-eGFP genome, most likely because of CRISPR-mediated homologous recombination.

Interestingly, recombination didn’t occur symmetrically on both side of the insertion site (**Fig. 2D**). The hCMV genome comprises two domains, each made of a unique segments (named UL for the longer segment, US for the shorter one) flanked by inverted repeats (*8, 17*). The two domains can be inverted relative to one another, forming four different linear configurations that can be isolated in equal amounts from viral particles (**Fig. 2E**). The gene drive cassette is located near one extremity of the UL fragment, and the *eGFP* and the BAC cassette are in the US segment. When mapping sequencing reads into the four linear genome configurations, we observed a similar pattern: SNPs around the gene drive cassette and in most of US segment originated predominantly from the donor TB40/E strain, but SNPs from the remaining of UL segment came predominantly from the Towne strain (**Fig. 2D, Fig. S5**). The enrichment for TB40/E SNPs in the US segment was unexpected. The genome of herpesviruses is linear, but circularizes after infection in the host cell nucleus. Homologous recombination might, therefore, occur in a circular genome, which exist in two configurations that both put the gene drive cassette in relative proximity to US segment (**Fig. 2E**). This and differences in the genome structure of the two viral strains may explain this complex recombination pattern.

Long-read sequencing further offered a unique opportunity to investigate recombination at the level of single viral genomes. Indeed, because Nanopore sequencing doesn’t involve PCR or any amplification step, each read originated from a distinct DNA molecule, and an individual sequencing read could capture the recombination history of one distinct viral genome. The hCMV genome is approximately 235 kb long, and we recovered 59 reads longer than 200 kb. After manually analyzing SNPs on mapped reads, we reconstructed the recombination history of 38 viral genomes (**Fig. 2F, Fig. S6**). Only three (8%) did not incorporate the gene drive cassette, and seven (18%) appeared to represent the pure TB40/E strain (the donor GD-mCherry viral genome). The remaining 28 sequences (74%) were recombination products of the Towne-eGFP and GD-mCherry genomes, and importantly, all included the gene drive cassette. Multiple genomes appeared to be exclusively from Towne origin, except for the incorporation of the gene drive cassette. On the other hand, some other sequences had incorporated major portions of both genomes. We also noted that most of US regions originated from TB40/E strain. These results demonstrated that the gene drive sequence can be efficiently and specifically transferred from the TB40/E strain to the Towne strain, creating new gene drive viruses. Homologous recombination appeared to be asymmetrically centered around the gene drive cassette, and created in some cases profound rearrangement between the two viral genomes. Thus, these data showed that gene drive virus could recombine with wildtype virus and incorporate the gene drive sequence into new virus.

We next aimed to show that the gene drive sequence could spread into a wildtype virus population. Fibroblasts were coinfected with wildtype Towne-eGFP (MOI=0.1) and decreasing amounts of GD-mCherry. Viral titers in the supernatant of coinfected cells were measured by plaque assay and allowed us to follow the evolution of the viral population over time (**Fig. 3A**). We observed that, independently of the starting proportion of GD-mCherry (50, 10, or 0.1% of the Towne-eGFP amount), the gene drive cassette efficiently invaded the wildtype population and incorporated into most of eGFP expressing viruses. Viruses expressing both eGFP and mCherry indeed represented new gene drive viruses that kept propagating the modification into the wildtype population. Importantly, we could barely observe the apparition of eGFP-mCherry recombinant virus when performing similar coinfection experiments with wildtype TB40/E or a *Cas9-*deleted version of the gene drive virus (GD-ΔCas9) (**Fig. 3B, Fig. S7**). This further confirmed that Cas9 is responsible for the drive of the recombination cassette into the wildtype population.

**Fig. 3.**
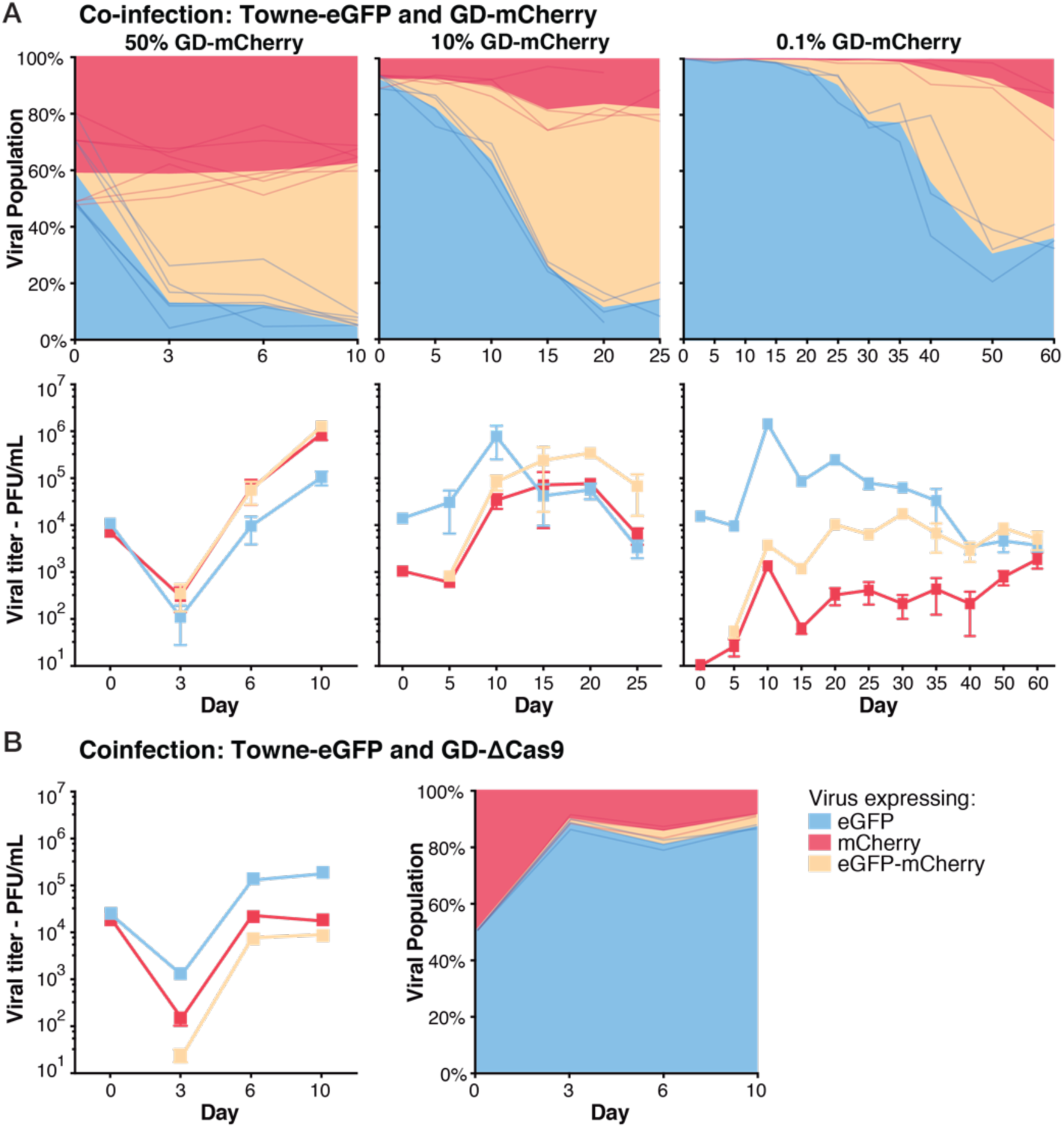
Gene drive sequences efficiently spread into the wildtype population. (**A**) Coinfection experiments between Towne-eGFP (MOI=0.1 at day 0) and different starting concentration of GD-mCherry. Viral titers (lower panels) and proportion (upper panels) over time of viruses expressing eGFP alone, mCherry alone, or both as measured by plaque assay. Left panel: 50% of GD-mCherry at Day 0, n=6; middle: 10% at D0, n=4; right: 0.1% at D0, n=3. (**B**) Viral titers and proportion of viruses after coinfection with equal amount of Towne-eGFP and GD-ΔCas9. n=3. Titers are expressed in PFU (plaque forming unit) per mL of supernatant. Error bars represent standard error of the mean (SEM) between biological replicates. In the panels representing the viral population, data show both the mean and the individual trajectory of biological replicates.

We showed the gene drive sequence can easily and efficiently spread into the wildtype population. However, a gene drive virus that replicates at levels similar to wildtype virus would have little therapeutic value. We wanted next to determine if a gene drive strategy could be used to limit or stop a viral infection. *UL23*, the viral gene targeted by our drive, was initially thought to be dispensable for hCMV replication in fibroblasts (*13, 18*). Serendipitously, it was later showed that UL23 is a tegument protein that blocks antiviral interferon-γ (IFN-γ) responses by interacting with human N-myc interactor (Nmi) protein (*19*). As a result, the growth of *UL23*-knockout viruses is severely inhibited in infected cells treated with IFN-γ. In our system, Cas9 target site is located immediately upstream of *UL23* coding sequence, and GD-mCherry viruses lack a *UL23* start codon (**Fig. S1A**). As predicted, we observed that the replication of GD-mCherry virus was strongly and significantly inhibited when cells were cultivated in presence of increasing concentrations of IFN-γ, with a 250- and 8000-fold titer reduction with 10 and 100 ng/mL of IFN-γ, respectively (**Fig. 4A, Fig. S8A** and **Fig. S8B** for thorough statistical analysis). Gene drive viruses are, therefore, strongly attenuated by IFN-γ antiviral response, while wildtype viruses are minimally affected.

**Fig. 4.**
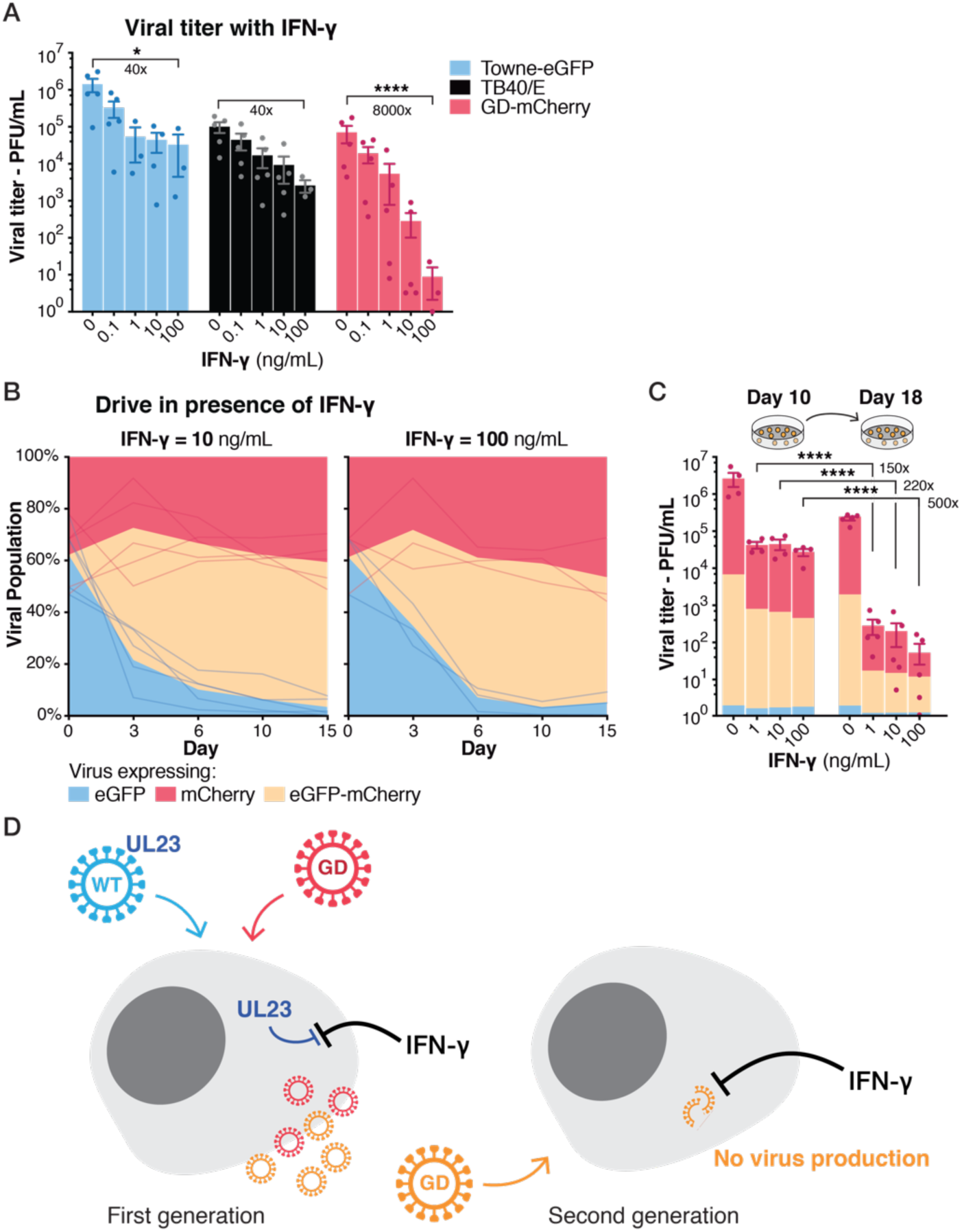
Spread of a defective gene drive virus. (**A**) Viral titers at day 10 in the presence of increasing concentrations of IFN-γ. n=3−5. (**B-C**) Drive with increasing concentrations of IFN-γ. n=3-5. (**C**) Cells were coinfected at Day 0 and supernatants were used to infect fresh cells at D10. Colors indicate the proportion of the different viruses relative to the height of the bar. n=4−5. (**D**) Model for the spread of defective gene drive viruses: Wildtype viruses express UL23 and block IFN-γ antiviral response while gene drive viruses are UL23-KO and are severely inhibited by IFN-γ. Upon coinfection, UL23 originating from the wildtype virus—either brought in with the incoming virion as a tegument protein or expressed early on—is sufficient to block the IFN-γ antiviral response. A first generation of new gene drive viruses can be created. These viruses are, however, UL23-KO and are severely inhib ited by IFN-γ when infecting new cells. Titers are expressed in PFU/ml. Error bars represent SEM between biological replicates.*, p-value < 0.05; ****, p < 0.0001; two-way ANOVA with Sidak’s multiple comparison test on log-transformed value.

The unique sensitivity of GD-mCherry viruses to different concentrations of IFN-γ also allowed us to test whether a severely defective virus could still efficiently drive into a wildtype population. We therefore repeated the coinfection experiment in the presence of increasing concentrations of IFN-γ (MOI=0.1). We observed that the efficiency of the drive was unaffected by IFN-γ and that most Towne-eGFP viruses efficiently incorporated the gene drive cassette (**Fig. 4B, Fig. S9A**). After 10 or 15 days of coinfection, most of the viral population had been converted and would have been expected to show high IFN-γ susceptibility. Unexpectedly, however, viral titers reached levels similar to wildtype viruses after 10 or 15 days, even at IFN-γ concentrations where GD-mCherry viruses are normally totally defective (**Fig. 4C, Fig. S9A**). This suggested that wildtype viruses were able to complement defective ones, blocking IFN-γ response and allowing the spread of new gene drive viruses in the context of a coinfection system (model in **Fig. 4D**).

However, even if the first generation of gene drive viruses appeared to be rescued by wildtype viruses, we hypothesized that subsequent generations would be severely inhibited by IFN-γ. To test this hypothesis, supernatants of coinfected cells at day 10 were used to infect fresh cells, treated with similar concentrations of IFN-γ. Viral titers of this second generation of viruses were severely affected by IFN-γ, with titers dropping 220- or 500-fold with 10 and 100 ng/mL of IFN-γ, respectively (**Fig. 4C**). Our gene drive against *UL23*, therefore, allowed us to drastically suppress viral infection and, in fact, represents an ideal scenario. In the presence of IFN-γ, gene drive viruses are almost non-infectious and cannot propagate by themselves. However, coinfection with a wildtype virus efficiently complemented the defective viruses and allowed their replication at high titer. As a consequence, the modification was able to spread efficiently into the viral population as long as wildtype viruses were present, but was unable to propagate further afterward (**Fig. 4D**). This represents an important example of how a viral gene drive could be used to cure or limit a viral infection.

By using hCMV as a model, we offer here a proof of concept that a gene drive can be successfully designed for herpesviruses. We show that modified viruses can spread in a viral population and replace their wildtype counterpart. We provide evidence that it is possible to design a drive that limits the infectivity of the virus. This represents a novel therapeutic strategy against herpesviruses and a new area of study. In the future, it will be interesting to determine if a similar viral gene drive can be designed against other herpesviruses or other DNA viruses and how the system can be adapted to function in an animal model and ultimately in human patients.

## Acknowledgments

We thank Edward Mocarski (Emory University) for providing viral stocks. MW thanks Rosalba Perrone and Kristal Fontaine for friendship and fruitful discussions.

## Author contributions

MW initiated and designed the study, performed all experiments and analyses. EV supervised and funded the project. MW and EV wrote the manuscript.

## Data and materials availability

All sequencing data have been deposited in the Short Read Archive with BioProject accession no. PRJNA545115. Plasmids, viruses and other reagents developed in this study are available upon request and subject to standard material transfer agreements with the Buck Institute.

## Material and Methods

### Cells and Viruses

Human foreskin fibroblast cells were obtained from the ATTC (#SCRC-1041) and cultured in DMEM (10-013-CV, Corning, Corning, NY, USA), supplemented with 10% FBS (Sigma-Aldrich, St-Louis, MO, USA) and 100 μ/mL penicillin/streptomycin (Corning). Cells were regularly tested negative for mycoplasma and used between passages 3 to 13.

hCMV TB40/E-Bac4 (*14*) and Towne-eGFP (T-BACwt) (*15*) were kindly provided by Edward Mocarski (Emory University, USA). To prepare viral stocks, cells were infected at low MOI (0.001–0.01) and kept in culture until 100% cytopathic effect was observed, usually after 10–15 days. Cells were then scrapped out of the plate and centrifugated together with the supernatant (10,000 rpm, 1 h, 4°C), resuspended in medium containing 5% milk, and sonicated to release cell-bound virions. Viral titers were assessed by plaque assay. Except when otherwise specified, subsequent infections were performed for 1 h at a MOI= 0.1, before replacing inoculum with fresh medium. Susceptibility to IFN-γ was assayed by virus growth in the presence of human recombinant IFN-γ (R&D, Minneapolis, MN, USA) after preincubation with IFN-γ for 2 h before infection.

Viral titers were assayed by plaque assay with 10-fold serial dilutions. 24-wells plates were inoculated for 1 h and overlaid with 0.25% agarose. After 7–10 days, eGFP or mCherry fluorescent plaques were manually counted using an inverted microscope. Every viral plaque was analyzed on both green and red channel. 5–100 plaques were counted per well, and each data-point was the average of 3–4 technical replicates (*i*.*e*., 3–4 different wells).

Coinfection experiments were performed by coinfecting with wildtype Towne-eGFP and gene drive viruses for 1 h, with a total MOI of 0.1–0.2. For time-course experiments over multiple weeks, supernatants were used to inoculate fresh cells for 1 h before changing media.

### Cloning and Generation of Gene Drive Viruses

A donor plasmid containing the gene drive cassette against UL23 (GD-mCherry) between homology arms was generated by serial modifications of pX330, a codon-optimized *SpCas9* (from Streptococus pyogenes) and chimeric gRNA expression plasmid developed by the Zhang lab (*20*) (Addgene #42230). All modifications were carried out by Gibson cloning (NEB, Ipswich, MA, USA), using PCR products from other plasmids or synthetized DNA fragments (GeneArt™ String™ fragments, ThermoFisher, USA). Briefly, a fragment with a SV40 polyA terminator, a SV40 promoter and an mCherry fluorescent reporter was inserted between *SpCas9* and betaGlobin polyA signal. The PciI-AgeI fragment upstream of *SpCas9* was removed and replaced by UL23 left homology arm (amplified from TB40/E-bac4). A fragment with UL23-5′ gRNA under a U6 promoter and UL23 right homology arm was finally inserted downstream of betaGlobin polyA signal between the NotI and Xmai restriction sites.

GD-ΔCas9 donor construct was subsequently generated by removing *SpCas9* by digestion and ligation. A donor construct to insert a SV40-driven mCherry reporter into the BAC cassette of hCMV TB40/E-bac4 was built similarly. Plasmids and sequences are available upon request.

To build gene drive viruses, 1.5 million fibroblast cells were transfected with the homologous recombination donor plasmid and an helper plasmid (Addgene #64221) (*21*). Transfection was performed by Nucleofection (Kit V4XP-2024, Lonza, Basel, Switzerland). 48 hours after transfection, cells were infected for 1 h with hCMV TB40/E-bac4 at a low MOI. After 7–10 days, viral plaques of mCherry-expressing cells were observed, suggesting successful integration of the gene drive sequence by homologous recombination (**Fig. S1B**). mCherry-expressing viral plaques were isolated and purified by several rounds of serial dilutions and plaque purification. Purity and absence of unmodified TB40/E viruses were assayed by PCR after DNA extraction (DNeasy kit, Quiagen). PCR and Sanger sequencing across homology arms and cut sites confirmed that mCherry-expressing viruses contained the full gene drive sequence (**Fig. S1B**). Viral stocks were produced as specified above and tittered by plaque assay.

### HIRT DNA Extraction, BAC Transformation and Analysis of Recombinant BAC Clones

hCMV episomal DNA was recovered from infected cells by HIRT DNA extraction (*16*) 48 h after infection at a high MOI. Infected cells (grown in 1–2 T175 plates) were scrapped-off, washed in PBS and resuspended in HIRT resuspension buffer (10 mM Tris-HCl pH 8.0, 10 mM EDTA in 100 μL for 1 million cells). An equal volume of HIRT lysis buffer (10 mM Tris-HCl pH 8.0, 10 mM EDTA, 1.2% SDS) was added and gently mixed. NaCl was added (1 M final concentration) before incubating overnight at 4°C. Supernatant was collected after centrifugation (13,000 rpm, 20 min, 4°C). Contaminating RNA was removed with RNAse A and hCMV DNA purified by double phenol-chloroform-isoamyl alcohol (25:24:1) extraction. DNA was precipitated with two volumes of pure ice-cold ethanol, washed with 70% ethanol, dried and resuspended in 10 mM Tris-HCl pH 8.0.

2–3 μL of recovered DNA was electroporated into NEB® 10-beta Electrocompetent cells (C320K, NEB) and plated on chloramphenicol LB plates. BAC DNA were purified using ZR BAC DNA Miniprep Kit (Zymo Research, Irvine, CA, USA).

Presence of *mCherry* or *GFP* on recombinant BAC clones was confirmed by PCR. Homology arms of mCherry-eGFP clones were analyzed by PCR and Sanger sequencing using tools available on Benchling.com. Primers are given in **Table S1**.

### Virion Purification and Oxford Nanopore Sequencing

Twelve large flasks were infected with recombinant virus and cultured until the monolayer reached 100% cytopathic effect. Cells debris was pelleted away by centrifugation (20 min, 3000 rpm, 4°C) and supernatants were recovered. Virions present in the supernatant were pelleted by ultracentrifugation (22 krm, 90 min, 4°C, Beckman-Coulter rotor SW28) on a 5-mL cushion of 30% sucrose. Supernatants were discarded, and the pellets containing virions were resuspended in PBS (1 h at room temperature) and pooled in a final volume of 500 μL.

DNase I (20U, 30 min, 37°C, NEB M0303S) and RNAse A (100 μg, 15 min, ThermoFisher EN0531) treatment removed contaminating human nucleic acids unprotected by the virus envelope and capsid. Viral envelopes were lysed and DNase I was inactivated with 5X lysis buffer (0.5 M Tris pH 8, 25 mM EDTA, 1% SDS, 1 M NaCl), and the lysate was incubated for 10 min. Incubation for 1 h at 55°C with 8U of proteinase K (NEB P8107S) finally disrupted viral capsids. To recover full-length hCMV genomes, subsequent pipetting and centrifugation steps were performed extremely carefully with wide-bore pipet tips. hCMV DNA was purified by double phenol-chloroform-isoamyl alcohol (25:24:1) extraction and centrifugation (6000 rpm, 3 min). DNA was precipitated for 1 h at -80°C with 1/20 volume of 5 M NaCl and 1.5 volume of cold ethanol, centrifugated (8000 rpm, 30 min, 4°C), washed with 70% ethanol, dried and resuspended in 10 mM Tris-HCl pH 8.0 for 24 h at 4°C without pipetting. DNA was quantified by Nanodrop.

Libraries were prepared using SQK-LSK109 ligation sequencing kit from Oxford Nanopore Technology (Oxford, UK), without any fragmentation steps, using wide-bore pipet tips and careful pipetting steps to minimize DNA shearing. Libraries for the first biological replicate (two technical replicates) were prepared, following the manufacturer instructions and using 2 μg of starting material. Lambda DNA control was added in the first technical replicate. In an attempt to maximize read length, the second biological replicate was prepared with 15 μg of DNA, omitting the first AMPure XP bead clean-up. Sequencing was performed on two FLO-MIN106-R9 Flow Cells on a MinION Mk1B device following manufacturer instructions.

### Oxford Nanopore Sequencing Analysis

Nanopore FAST5 raw data was converted into FASTQ files using Albacore v2.3.3 basecalling, and runs statistics were obtained using Nanoplot (https://github.com/wdecoster/NanoPlot) (*22*). Adaptors were removed with Porechop (https://github.com/rrwick/Porechop). Reads with quality Q > 6 were filtered using Nanofilt (https://github.com/wdecoster/nanofilt) (*22*) and Lambda DNA reads excluded with NanoLyse (https://github.com/wdecoster/nanolyse) (*22*). Technical replicates were merged for the rest of the analysis. Reads lengths were filtered with Nanofilt.

Reference sequences for Towne-eGFP (GenBank KF493877) and TB40/E-Bac4 (GenBank EF999921) were downloaded. We first inserted the gene drive cassette into Towne-eGFP and TB40/E reference sequences. Reads were then mapped on a composite human hg38-Towne genome using Minimap2 (https://github.com/lh3/minimap2) (*23*) and mapping statistics (**Fig. S4**) were obtained with samtools (*24*) after filtering of secondary and supplementary reads (`samtools -F 2048 -F 256`).

To create of map of SNPs between Towne and TB40 strains, we created a FASTA file composed of multiple copies of the two genomes. Using Minimap2, this fasta file was mapped onto Towne-eGFP reference. SNPs were then called using bcftools mpileup and bcftools call, generating a BCF file with the SNPs coordinates:

~~~
minimap2 -a Ref_Towne.fasta multiple_Towne-TB40.fasta -cs -N 0 -2 | samtools view -S -b| samtools sort -o mapped.bam
bcftools mpileup -Ou -f Ref_Towne.fasta mapped.bam | bcftools call -mv -Ob -o map_snp.bcf
~~~

hCMV genome exist in four different configurations depending on the respective orientation of UL and US segments. A Towne-eGFP composite genome composed of Towne sequences in the four configurations was finally created, and a complete map of SNPs in the four configurations was also generated.

After comparing different mappers, we mapped sequencing reads (length > 10kb) on the composite Towne genome using Graphmap (https://github.com/isovic/graphmap) (*25*), keeping only the best mapped reads (default, returning uniquely mapped read) or allowing multiple read mapping (option -Z). Proportion of variants for each SNP coordinate was then calculated using Nanopolish (https://github.com/jts/nanopolish) (*26*). Nanopolish was run successively on the 4 subgenomes as follow, using the SNP map generated above (example for Towne in the SS: Sense-Sense configuration):

~~~
nanopolish variants --reads data.fastq.gz --bam Mapped_with_graphmap.bam -- genome Reference_Towne_4config.fasta -p 2 -w Towne_config_SS:1-239862 -m 0.15 -x 2000 -c map_snp_4config.vcf -o Call_SS.vcf
~~~

Read coverage and Support fractions were then extracted from the VCF files and plots were generated using R. Of note, for multiple mapping reads, we had to artificially increase the mapQ fields of SAM/BAM files by 20, because Nanopolish automatically discard such reads.

Finally, to reconstruct the recombination history of individual genomes, the longest reads (>200 kb) were mapped on the composite Towne genomes using Graphmap and visualized with IGV (*27*). The recombination map of each individual reads was then reconstructed manually using the SNP map.

The reference genome of GD-mCherry virus was inferred from reads that contained no Towne fragments. Importantly, we detected two large deletions in GD-mCherry not present in the original TB40E-bac4 genome: One 5.6-kb deletion in UL ranging from *RL12* to *UL8* genes and, therefore, including the frequently mutated *RL13* gene, and a second 4.6-kb deletion in US from *US15* to *US19*.

### Statistical analysis of plaque assay data

Plaque assay data didn’t appear to satisfy the normality condition required for parametric tests. Due to small sample sizes, normality and lognormality tests could however not be performed. We therefore chose to run in parallel both parametric tests on log-transformed data, and non-parametric test on untransformed data, and reported results when both type of tests gave significant values. For groups analysis, we performed two-way ANOVA with Sidak’s multiple comparison test on log-transformed data, and Krustal-Wallis test with Dunn’s multiple comparison test on untransformed data.

Analysis were run using GraphPad Prism version 8.1.1 for macOS (GraphPad Software, San Diego, California USA, www.graphpad.com)

**Table S1:**
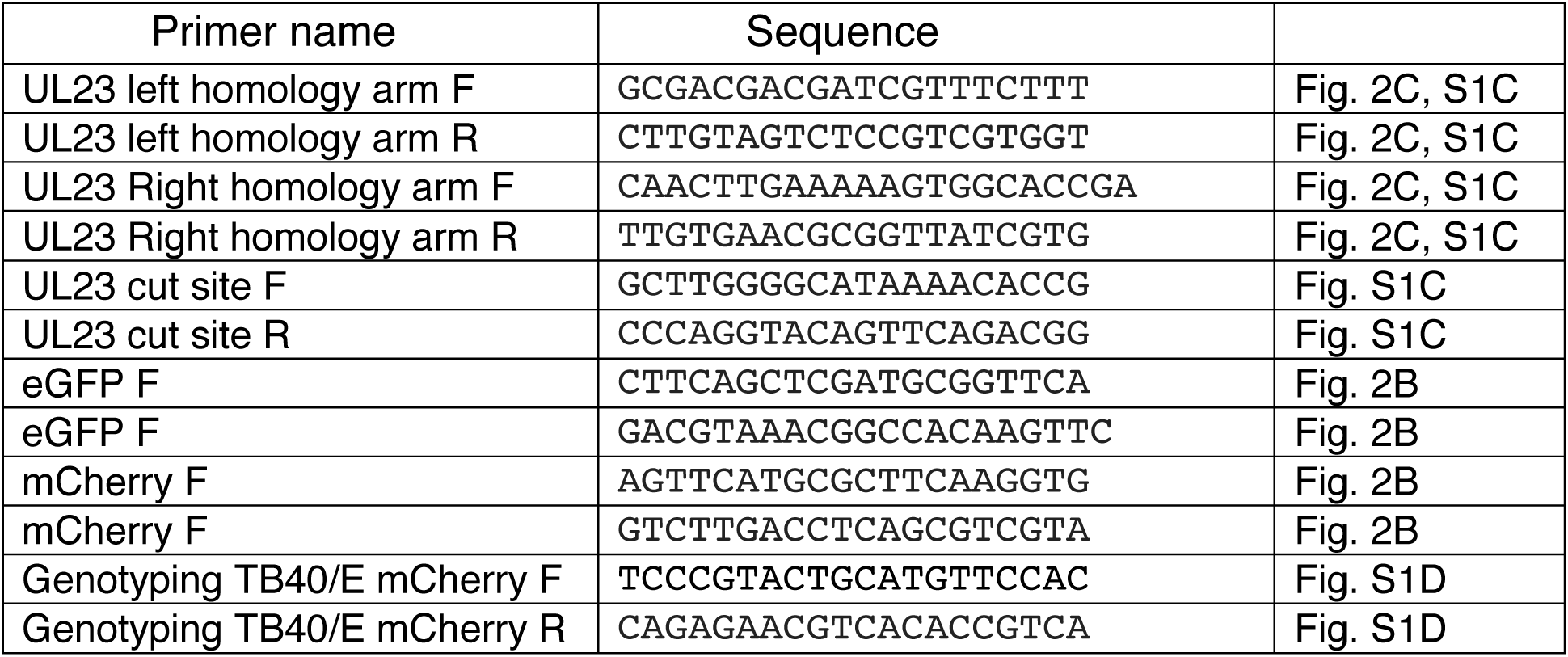
Primers.

**Fig. S1.**
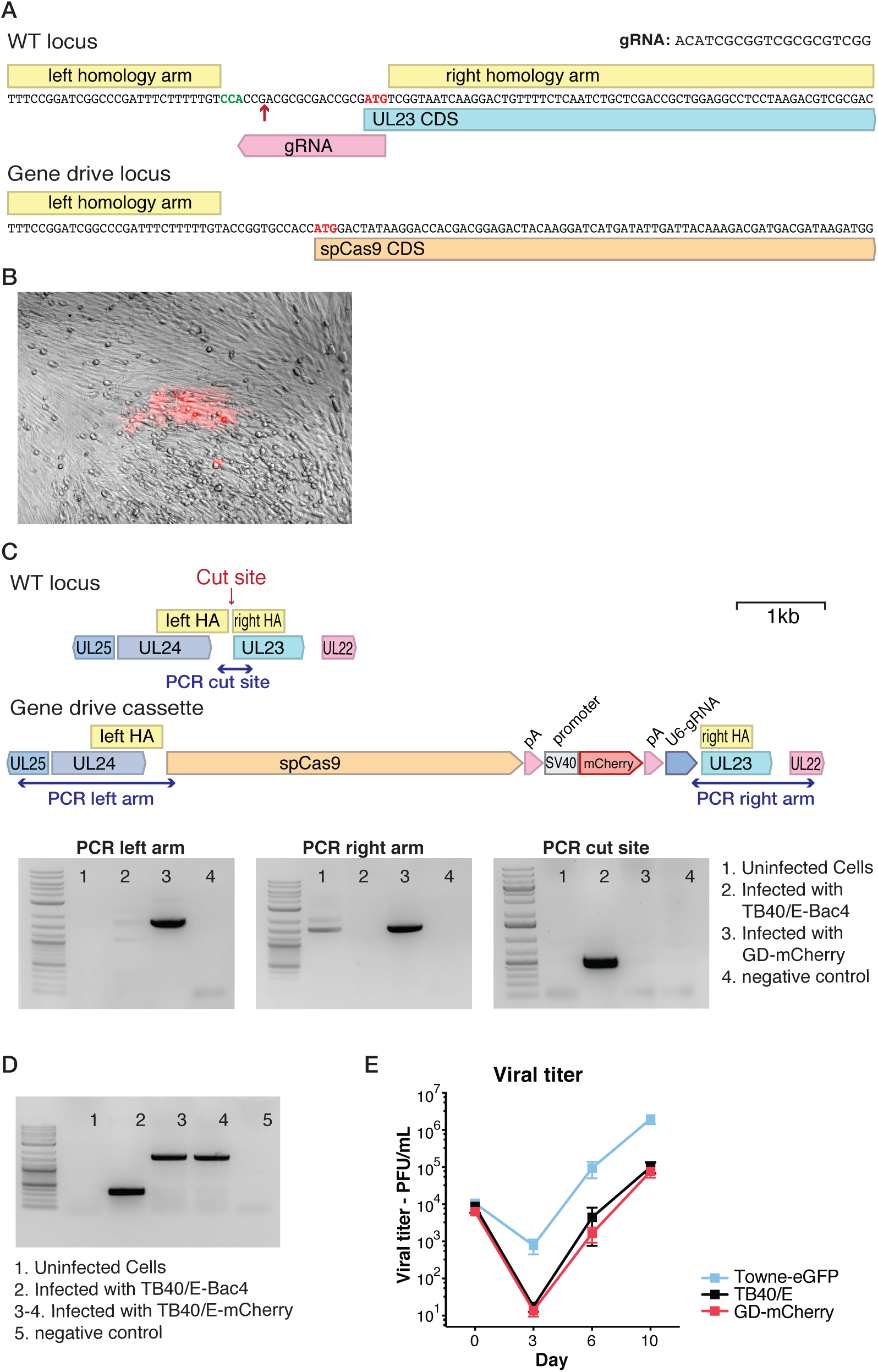
Generation of gene drive viruses. (**A**) UL23 CRISPR cutting site and sequence of wildtype (WT) and modified viruses. (**B**) Representative image of a mCherry-expressing viral plaque spreading into human fibroblasts, approximately 10 days after transfection with the gene drive donor plasmid and infection with hCMV (TB40/E strain). (**C**) Genotyping PCR of pure GD-mCherry population and primer localization. (**D**) Genotyping PCR of pure TB40/E-mCherry population. (**E**) Viral titers were measured by plaque assay over time. Titers are expressed in PFU (plaque forming unit) per mL of supernatant. Error bars represent standard error of the mean (SEM) between biological replicates. n=4–6.

**Fig. S2.**
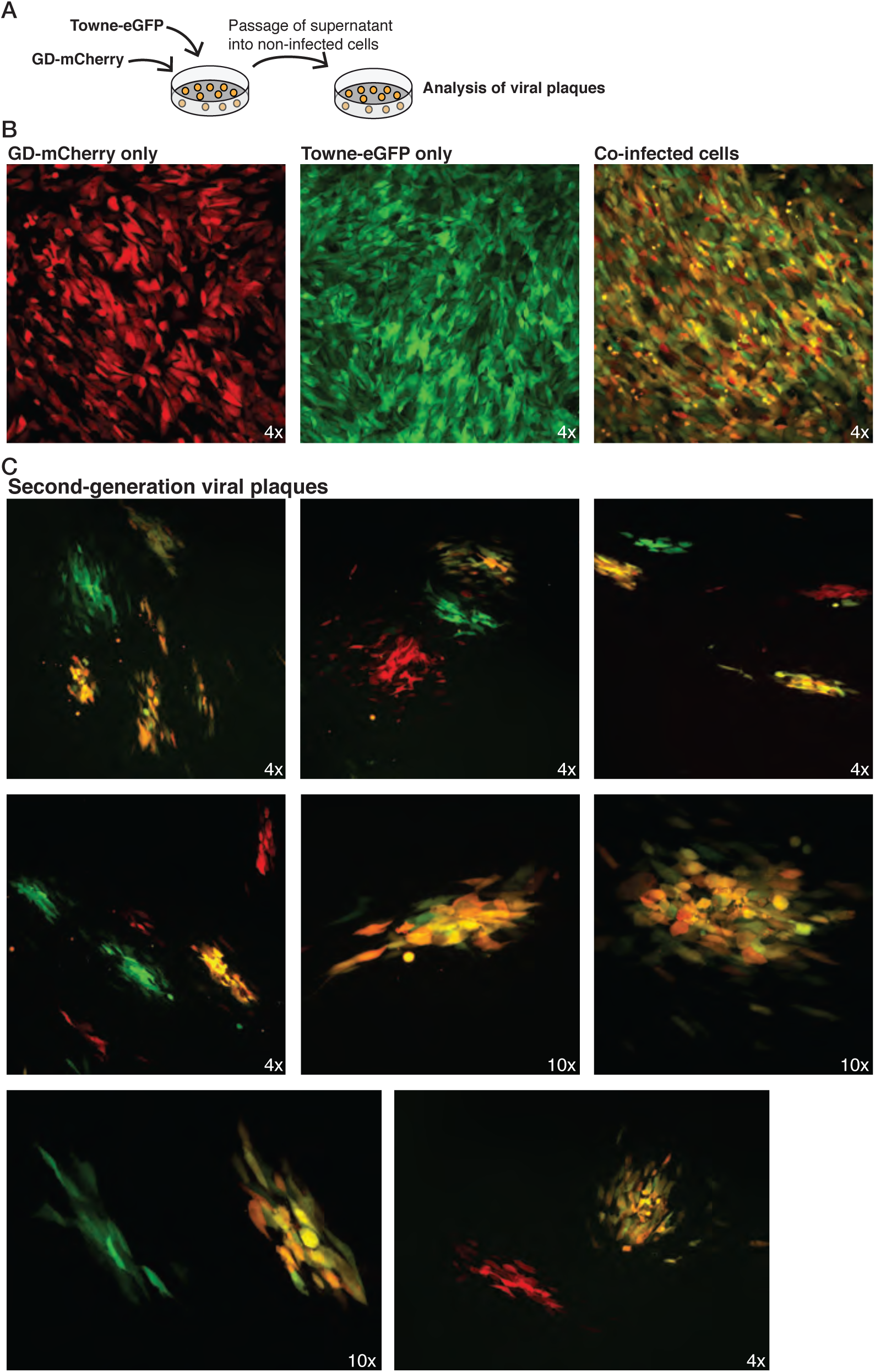
Recombinant viral plaques. (**A**) Experimental scheme: fibroblasts were coinfected with Towne-eGFP and GD-mCherry, and supernatants used to infect fresh cells. (**B**) Images of cells infected with Towne-eGFP, GD-mCherry, or both. (**C**) Images of second generation recombinant viral plaques.

**Fig. S3.**
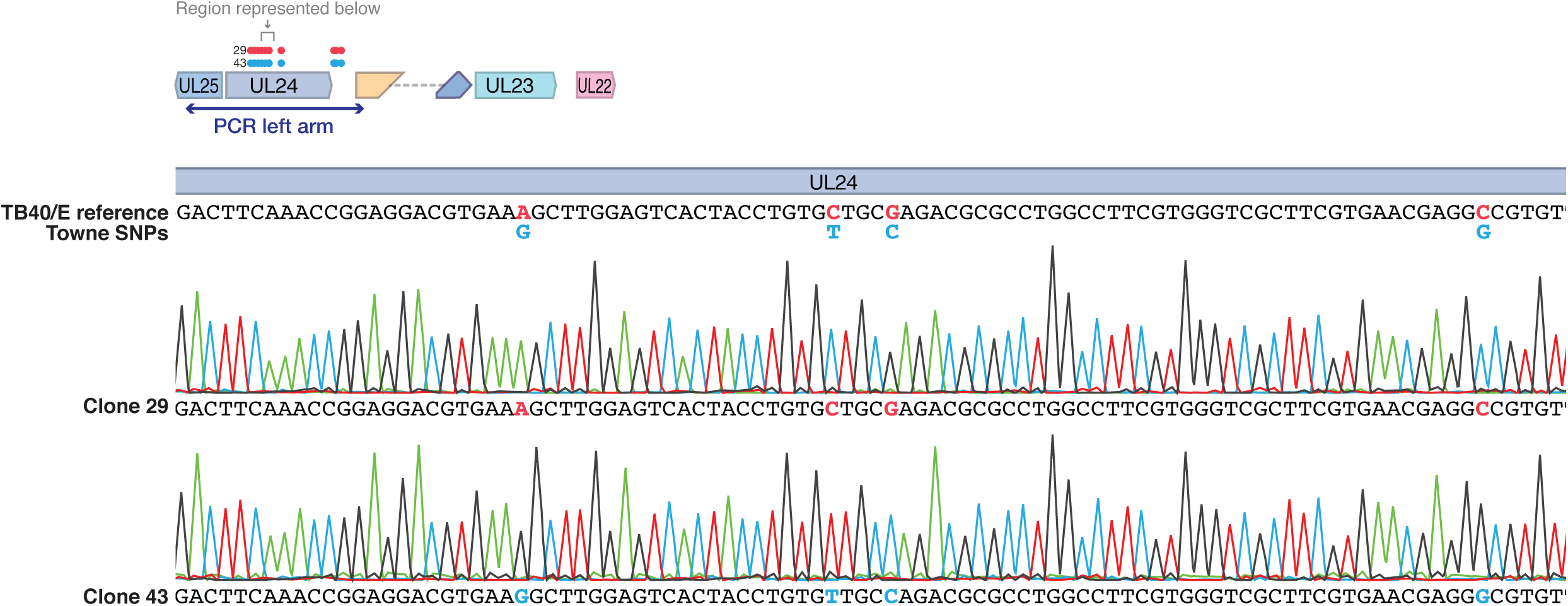
Sanger sequencing of homology arms. Example of Sanger sequencing of the left homology arm of two BAC clones. Clone 29 harbor SNPs from TB40/E strain, and clone 43 has SNPs from Towne strain.

**Fig. S4.**
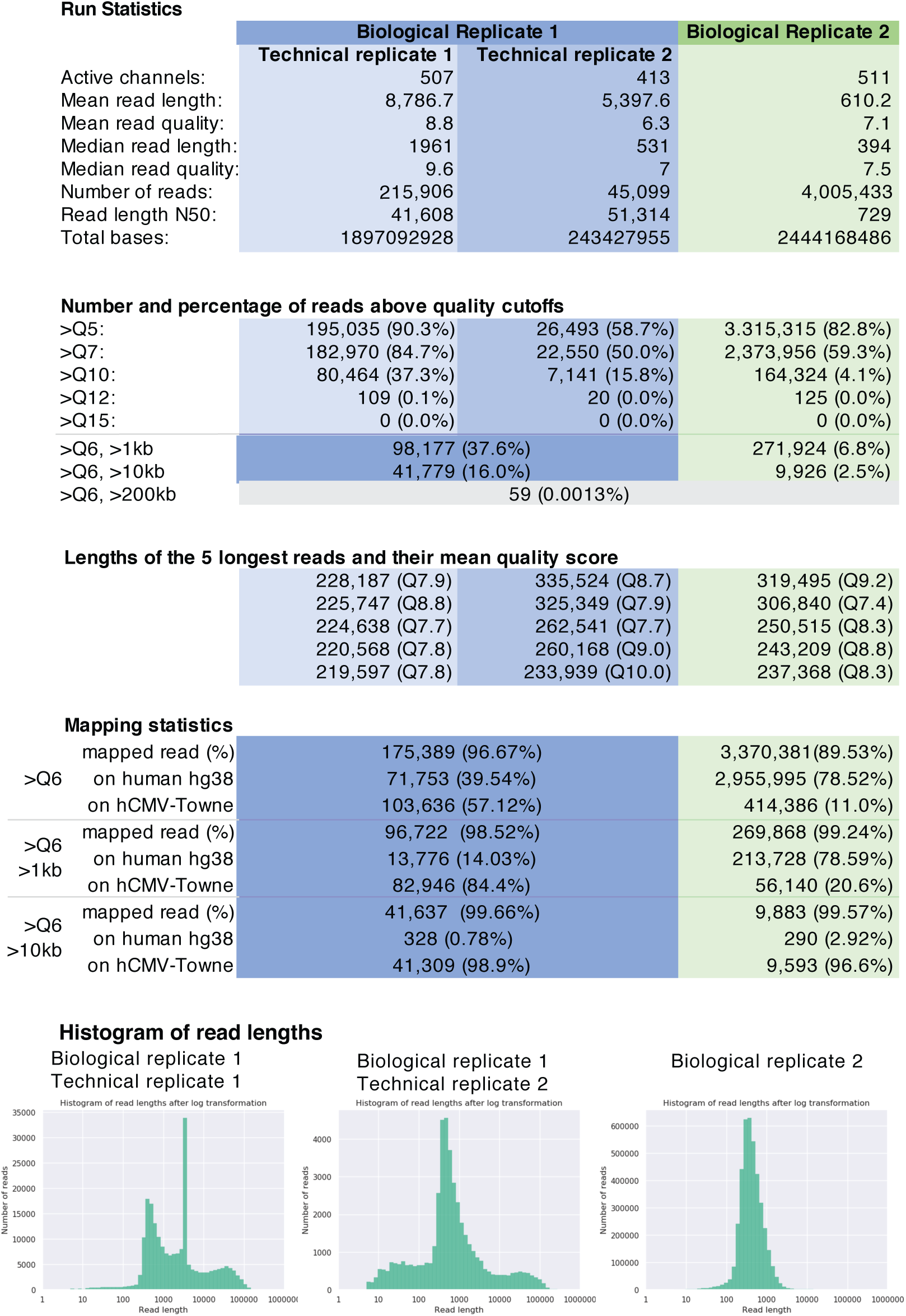
Oxford Nanopore sequencing statistics. Linear viral DNA was extracted from virions and subjected to long-read sequencing using Oxford Nanopore sequencing. The first biological replicate was sequenced in two technical replicates.

**Fig. S5.**
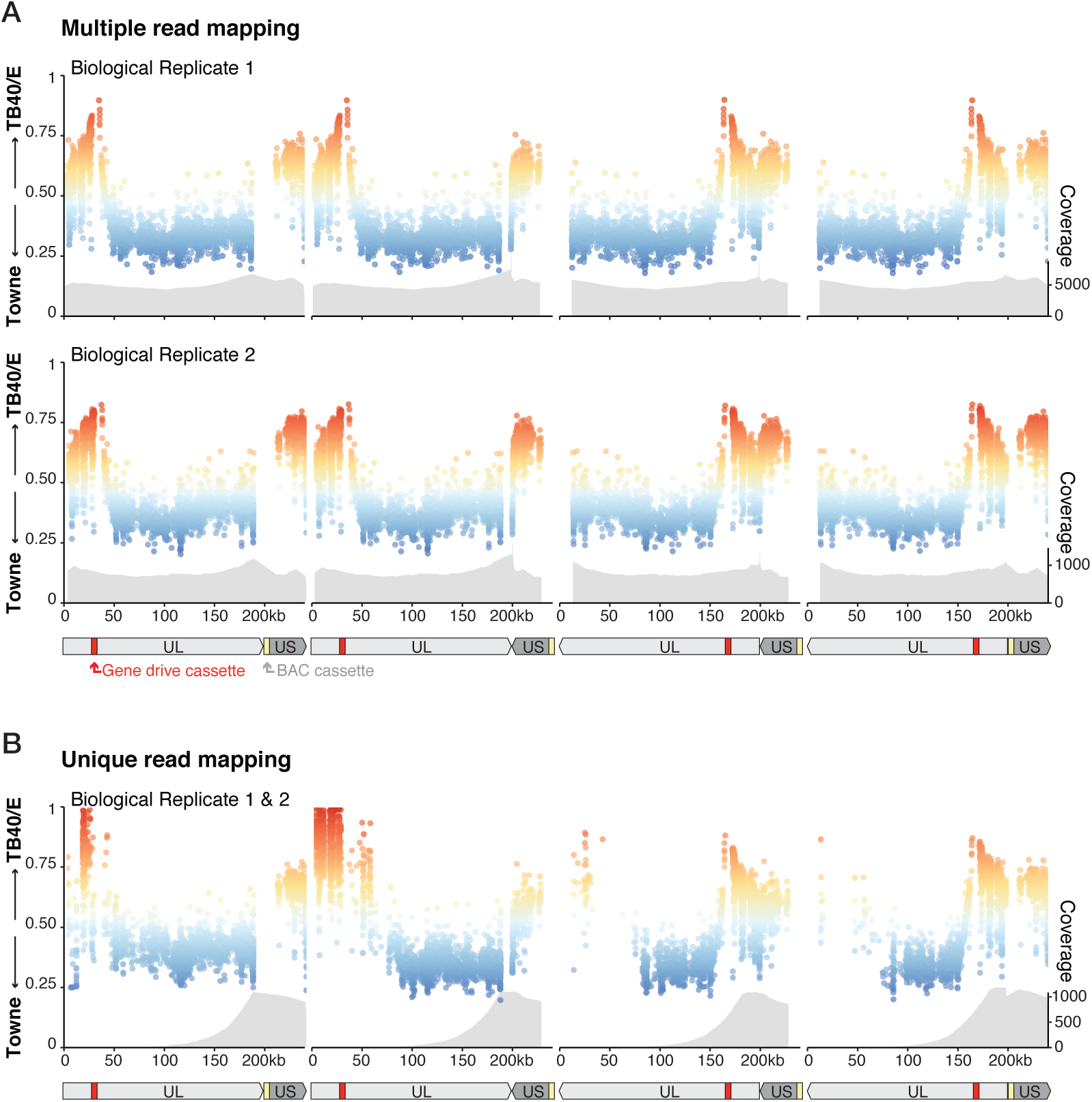
SNP proportion in the four genome orientations. (**A**) Fraction of SNPs of Towne or TB40/E origin in the four different genome configurations. Each dot represents an individual SNP. Coverage gives the number of reads. Reads mapping ambiguously into one or multiple genome configurations. (**B**) Same as (A) but with reads mapping unambiguously into a unique genome configuration.

**Fig. S6.**
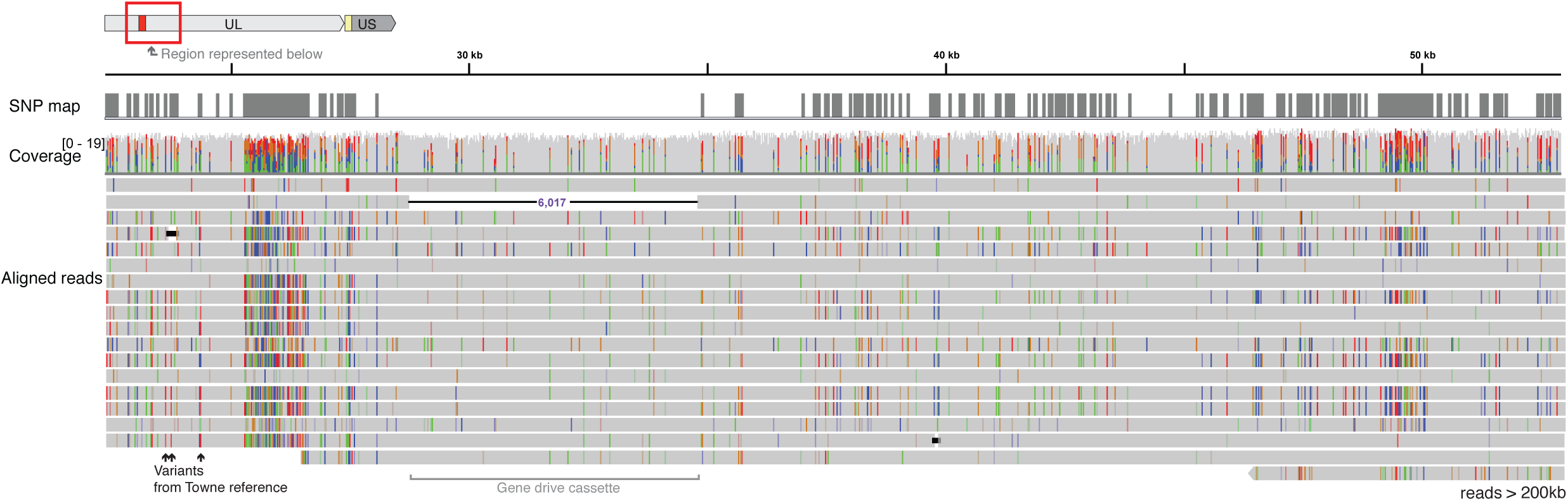
Reconstruction of recombination history from long reads. IGV screenshot showing individual long reads (>200 kb) mapped into Towne reference sequence. Reads are extremely noisy with numerous errors, but clusters of variants matching the SNP map allowed us to reconstruct the strain of origin.

**Fig. S7.**
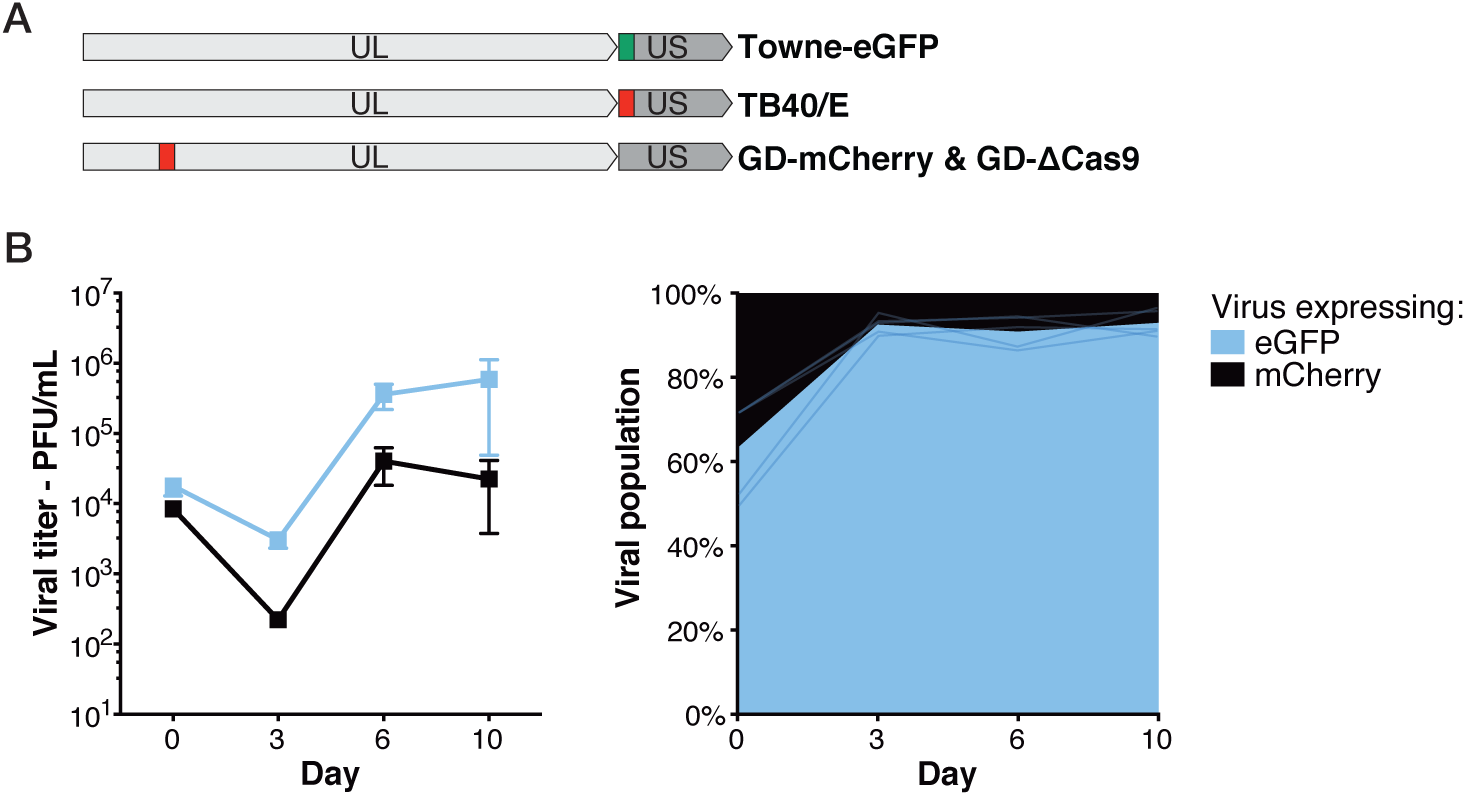
Coinfection with Towne-eGFP and TB40/E. (**A**) Localizations of mCherry and eGFP cassette on hCMV genomes. (**B**) Viral titer and proportion of viruses expressing eGFP alone, mCherry alone, or both, after coinfection with equal amount of Towne-eGFP and TB40/E. n=5. Titers are expressed in PFU/mL. Error bars represent SEM between biological replicates.

**Fig. S8.**
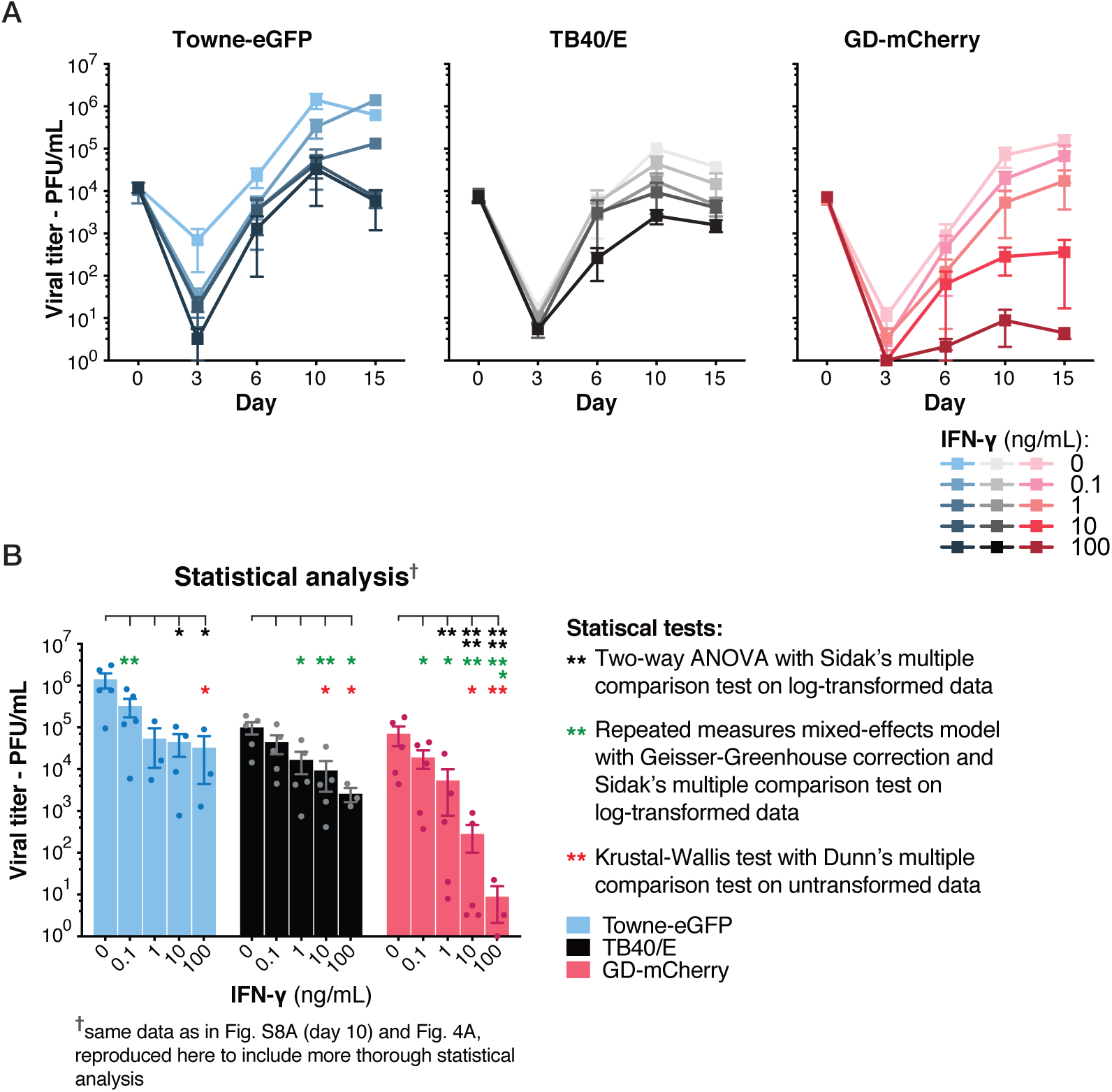
Viral titer in presence of IFN-γ. (**A**) Viral titer of single infections, in the presence of IFN-γ, measured by plaque assay. n=3–5. Data for D10 are also shown in Fig. 4A. and Fig. S8B. (**B**) Viral titer at D10 in presence of increasing concentration of IFN-γ. n=3–5. Same figure as Fig. 4B with thorough statistical analysis. Titers are expressed in PFU/mL. Error bars represent SEM between biological replicates. *, p-value < 0.05; **, p < 0.01; ***, p < 0.001; ****, p < 0.0001. Of note, two-way ANOVA or other parametric tests on untransformed data gave no significant results.

**Fig. S9.**
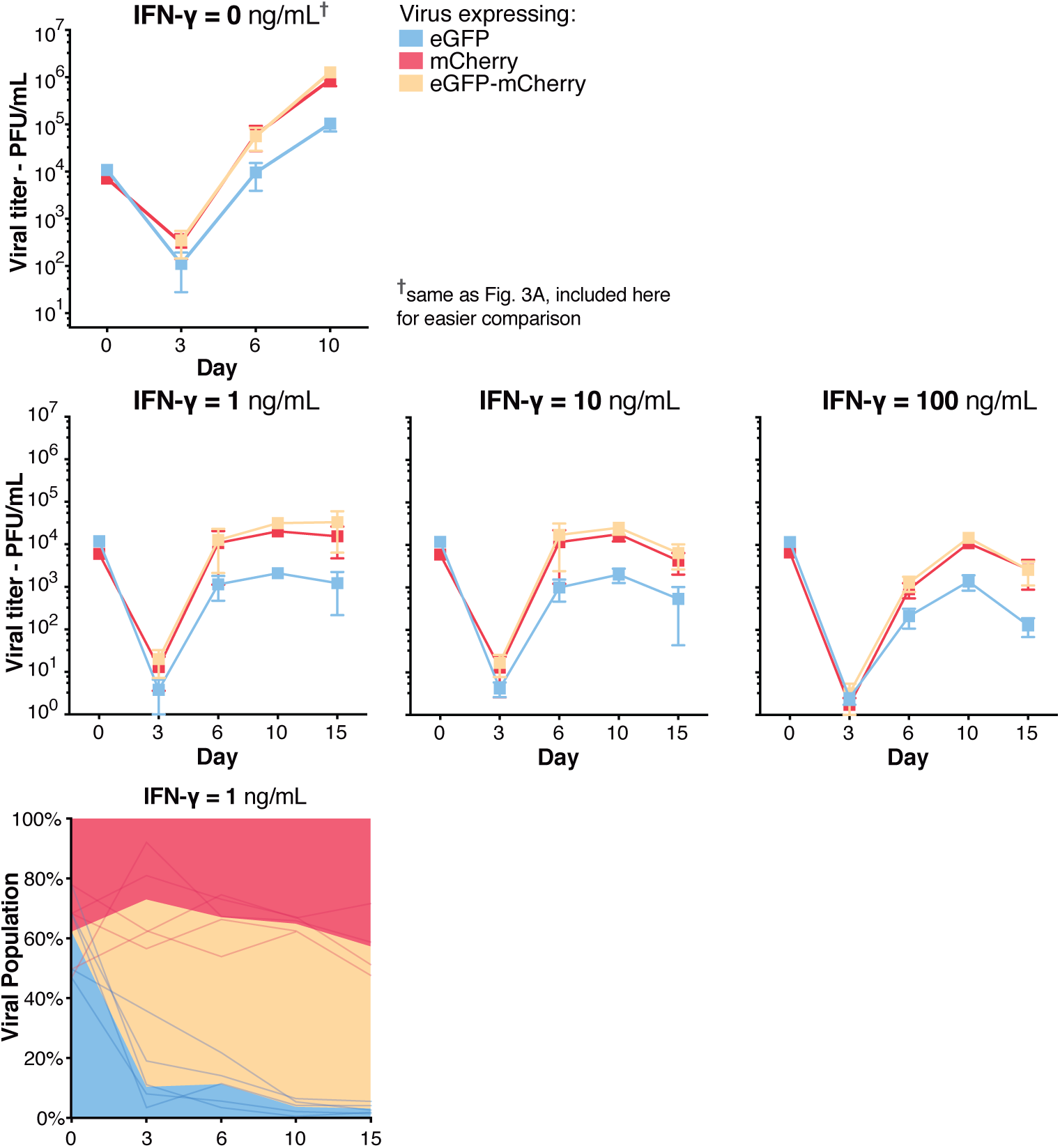
Gene drive in the presence of IFN-γ. Viral titer and proportion of viruses expressing eGFP alone, mCherry alone, or both, after coinfection with equal amounts of Towne-eGFP and GD-mCherry, in presence of increasing concentrations of IFN-γ. Same experimental data as shown in Fig. 4B-C. n=3–5. Titers are expressed in PFU/mL. Error bars represent SEM between biological replicates.

